# BAllC and BAllCools: Efficient Formatting and Operating for Single-Cell DNA Methylation Data

**DOI:** 10.1101/2023.09.22.559047

**Authors:** Wei Tian, Wubin Ding, Jiawei Shen, Daofeng Li, Ting Wang, Joseph R. Ecker

**Affiliations:** Genomic Analysis Laboratory, The Salk Institute for Biological Studies, La Jolla, CA 92037, USA; Department of Genetics, The Edison Family Center for Genome Sciences & Systems Biology, Washington University School of Medicine, St. Louis, MO 63110, USA; McDonnell Genome Institute, Washington University School of Medicine, St. Louis, MO 63108, USA; Howard Hughes Medical Institute, The Salk Institute for Biological Studies, La Jolla, CA 92037, USA

## Abstract

**Motivation:** With single-cell DNA methylation studies yielding vast datasets, existing data formats struggle with the unique challenges of storage and efficient operations, highlighting a need for improved solutions.

**Results:** BAllC (Binary All Cytosines) emerges as a tailored binary format for methylation data, addressing these challenges. BAllCools, its complementary software toolkit, enhances parsing, indexing, and querying capabilities, promising superior operational speeds and reduced storage needs.

**Availability:** https://github.com/jksr/ballcools

**Supplementary information:** Supplementary data are available at Bioinformatics online.

## 1 Introduction

The advent of single-cell DNA methylation (scDNAm) research has significantly enhanced our grasp of epigenetic intricacies, bringing to light the heterogeneities existing across individual cells – heterogeneities that are often masked in bulk assays (Luo *et al*., 2017; Liu *et al*., 2021; Tian *et al*., 2023). However, the granularity and scope of such investigations yield immense datasets, often spanning tens to hundreds of terabytes, presenting storage and processing challenges. For instance, Neuroscience Multi-Omic Data Archive (NeMO Archive), a prominent omic database for brain science, harbors over four hundred terabytes of scDNAm data, occupying more than half of its total storage capacity (as of Mar 22, 2024). Moreover, it has been revealed that >40% of the storage space in the NCBI Gene Expression Omnibus (GEO) is allocated to scDNAm data (private conversations). Such substantial storage demands for scDNAm data have become significant technical obstacles and financial burdens for both researchers and databanks. While various specialized formats have been developed for DNA methylation profiling (Argelaguet *et al*., 2019; Li *et al*., 2021; Ding *et al*., 2022; Loyfer *et al*., 2023), many fail to adequately address the distinct sparsity of scDNAm, encompass both CpG and non-CpG methylation contexts, or achieve the optimal operational speed and storage efficiency. Recognizing the pressing need for a more efficient alternative, we embarked on the development of BAllC (<u>B</u>inary <u>All C</u>ytosines), a tailored binary format for single-cell DNA methylation data. Accompanied by its software toolkit, BAllCools, this system aims to counter the limitations of existing formats, offering researchers a streamlined approach for the next era of epigenetic exploration.

## 2 Methods

### 2.1 The BAllC format

The BAllC format comprises a header and a methylation record section (Figure 1). The header identifies the BAllC version, provides genome reference information, and accommodates user notes. The methylation record section contains genome coordinates and both methylated and total counts for each cytosine. To leverage the inherent bound for non-duplicated counts of individual cytosines in single-cell as well as accommodating the frequent need to merge single-cell data to pseudo-bulk levels, the BAllC format employs different data types for single-cell and bulk files (uint8 and uint16, respectively; Figure 1). This strategy optimizes storage efficiency while maintaining data integrity. A supplementary TAB-delimited CMeta file is used to store the positions and contexts of each cytosine in the genome (Table S1). Only one CMeta file is required for multiple BAllC files sharing the same genome reference, minimizing data redundancy. Both BAllC and CMeta files Tian et al. employ the blocked compressed format (BGZF (Bonfield *et al*., 2021)) and can be indexed for rapid data retrieval.

### 2.2 BAllCools software package

BAllCools is a C/C++ software package designed for operations on BAllC files. It supports format conversion between BAllC and the AllC text-based format (Schultz *et al*., 2015), provides indexing and querying capabilities for BAllC files, and offers efficient methylation information retrieval based on user-defined genome range(s), strandness, or cytosine context. Notably, BAllCools can merge multiple BAllC files rapidly, facilitating analysis of methylomes at cluster or cell-type levels. Additionally, Python and Javascript wrappers are also available to enhance flexibility in utilizing the BAllC format.

**Figure 1.**
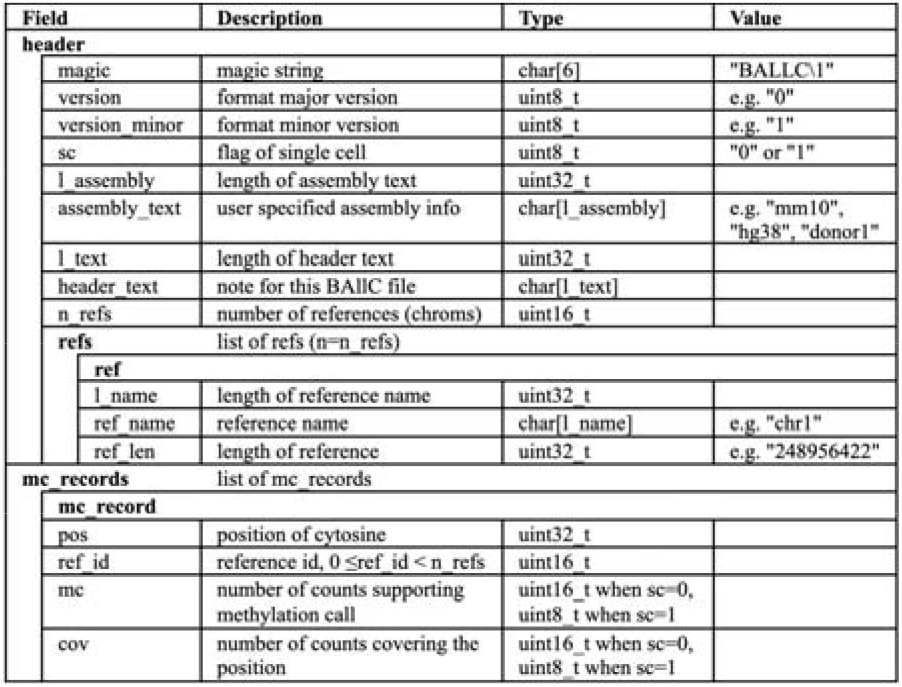
Specification of the BAllC format.

## 3 Results

We evaluated the efficiency of BAllC/BAllCools using large public single-cell DNA methylation datasets (Liu *et al*., 2021; Tian *et al*., 2023) and benchmarked them against the text-based AllC format (Schultz *et al*., 2015) and the associated toolkit AllCools (Liu *et al*., 2021), both of which are extensively used in studies of single cell DNA methylation.

### 3.1 Storage efficiency

Due to its binary nature and minimized redundancy, the BAllC format significantly optimizes storage in comparison to text-based formats. Specifically, the BAllC format achieved a >94% reduction in storage compared to the uncompressed text-based format. When compared to the compressed text-based format (BGZF format), the bulk mode BAllC format achieved approximately 50.4% reduction in storage, while the single-cell BAllC mode format achieved about 55.2% reduction.

### 3.2 Operational speed

Merging multiple single-cell methylomes into pseudo-bulk level is one of the most frequent operations in processing scDNAm datasets, permitting further analyses like differential methylation calling and visualization on genome browsers. Merging with BAllCools drastically cuts down the time required in comparison to text-based alternatives (Figure S1). For instance, merging a typical set of 500 scDNAm data takes only about 1.8 hours using one CPU core (2.80GHz), resulting in over 47x time savings compared to the text-based alternative.

### 3.3 Flexible data query and visualization

BAllCools extends its functionality beyond mere retrieval of DNA methylation records for specific genome coordinates. It offers users the capability to refine their queries by considering strandness and cytosine context. Users can retrieve methylation statuses of only CpG sites or cytosines within contexts of particular interest (Figure S2). These flexibilities are often unattainable with the text-based format and its associated tools. BAllCools also facilitates various operations on the queried regions, including aggregation of methylated and unmethylated counts, cytosine count tallying, and average methylation level calculations. These operations furnish essential metrics at the genome region level. The BAllC format file can also be visualized in the WashU Epigenome Browser (Li *et al*., 2019) for checking methylation status (Supplementary Data).

## 4 Conclusions

The increasing volume of single-cell DNA methylation data provides valuable resources for understanding gene regulation and heterogeneity. Despite the availability of various well-established and effective analysis tools such as DSS, methylKit, MethPipe, AllCools, Methylpy, etc. (Feng *et al*., 2014; Akalin *et al*., 2012; Song *et al*., 2013; Liu *et al*., 2021; Schultz *et al*., 2015), the challenge of storage remains unaddressed. In response to this challenge, we introduce the BAllC format as an efficient storage solution for scDNAm data. This binary format, in conjunction with the efficient toolkit BAllCools, represents a significant advancement in handling scDNAm datasets, ensuring optimal performance in terms of storage, speed, and data retrieval.

## Acknowledgements

We thank Jingtian Zhou for assistance in testing and for suggestive comments. We are very grateful to members of the Ecker group for their feedback and discussion.

