## Supplementary material for "BAllC and BAllCools: Efficient Formatting and Operating for Single-Cell DNA Methylation Data": ballc Supplemental material.docx.pdf

### Tutorial: Visualizing BALLC format files on the WashU Epigenome Browser

This tutorial shows you how to visualize the methylation data encoded in the BALLC format on the WashU Epigenome Browser.

1. Open the Browser at https://epigenomegateway.wustl.edu/browser/, here we chose the human hg38 as our genome and kept only the refGene track.

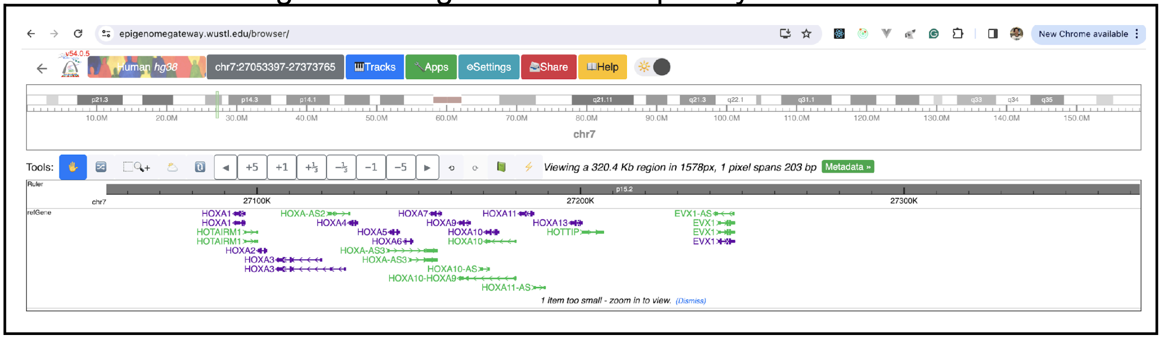


2. Add a BALLC file hosted from your web server using the Remote Tracks open dialog:

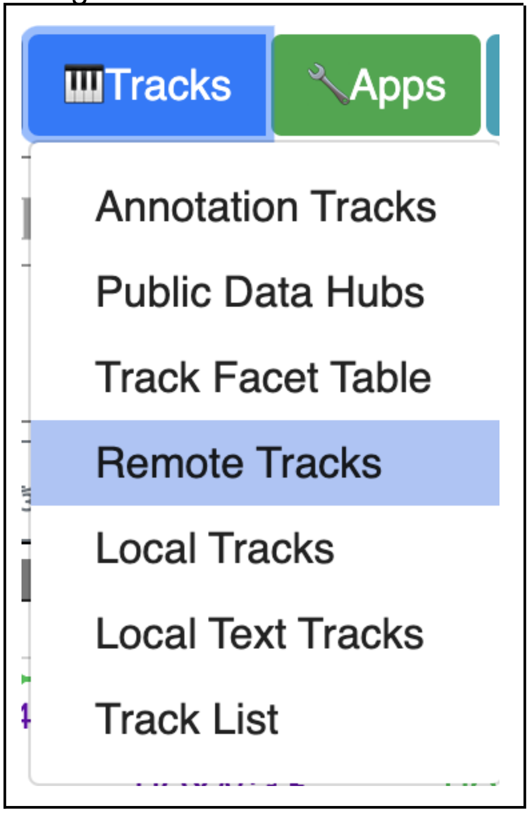

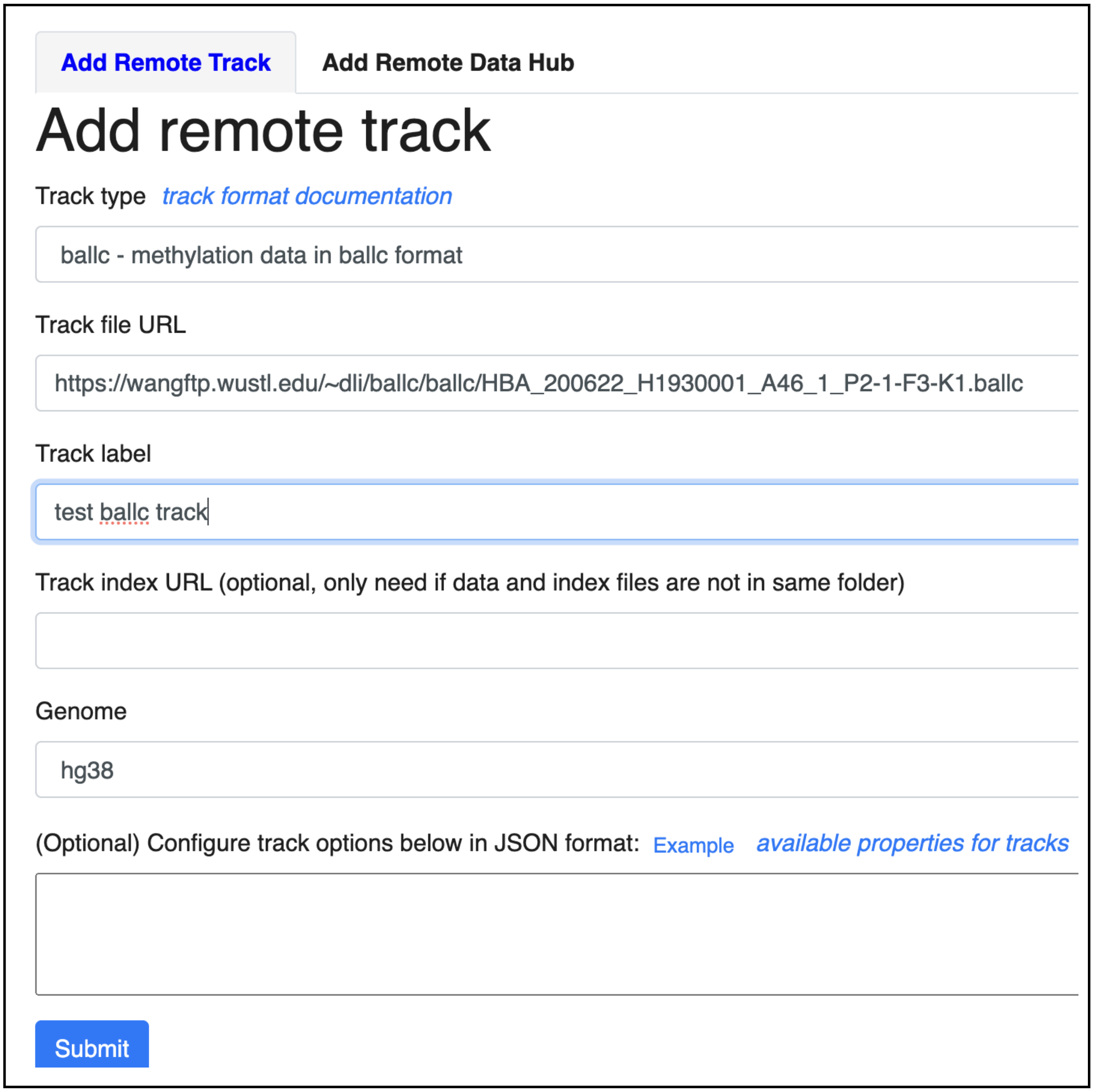


3. Click the Submit button, the BALLC file will be display in the Browser:

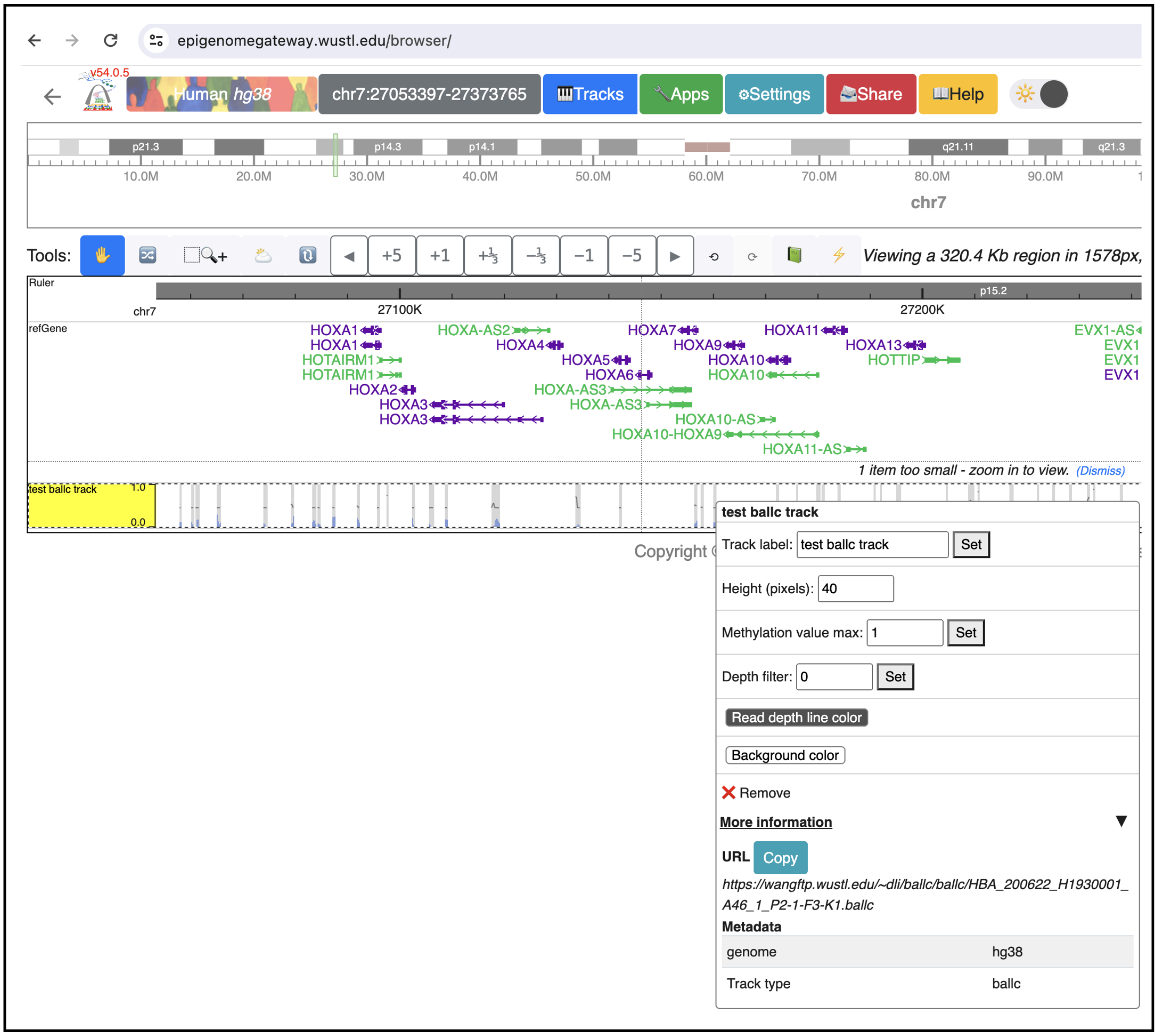


Each bar in the track indicates methylation signal in the region summarized from the CpG sites in it, blue bar indicates methylation percentage from 0 to 1, black line represents read depth/coverage.

If we zoom into base pair level view:

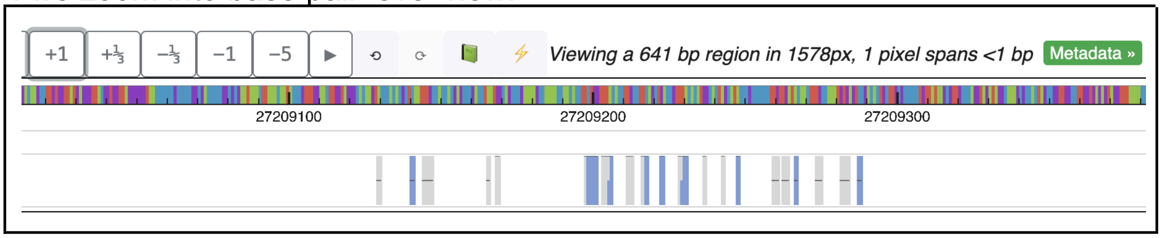


Each bar represents methylation status of a C base in the genome; a gray bar without blue color means that C is fully unmethylated, whereas a full blue bar means that C base is fully methylated. Black line indicates read depth.
