## Supplementary material for "BAllC and BAllCools: Efficient Formatting and Operating for Single-Cell DNA Methylation Data": figS2.context-query.pdf

A

```
~$ ballcools query example.ballc chr5:948645-948696
--cmetapath example.cmeta.gz
chr5    948645  +       CTG     0       1       1
chr5    948651  +       CGT     1       1       1
chr5    948660  +       CGG     1       1       1
chr5    948667  +       CTG     0       1       1
chr5    948671  +       CGC     1       1       1
chr5    948673  +       CGT     1       1       1
chr5    948677  +       CTA     0       1       1
chr5    948686  +       CCA     0       1       1
chr5    948687  +       CAG     0       1       1
chr5    948690  +       CAT     0       1       1
chr5    948693  +       CTT     0       1       1
chr5    948696  +       CTG     0       1       1
```

B

```
~$ ballcools query example.ballc chr5:948645-948696
--cmetapath example.cmeta.gz --c_context CGN
chr5    948651  +       CGT     1       1       1
chr5    948660  +       CGG     1       1       1
chr5    948671  +       CGC     1       1       1
chr5    948673  +       CGT     1       1       1
```

C

```
~$ ballcools query example.ballc chr5:948645-948696
--cmetapath example.cmeta.gz --c_context CTG
chr5    948645  +       CTG     0       1       1
chr5    948667  +       CTG     0       1       1
chr5    948696  +       CTG     0       1       1
```

chr pos strand context mc cov
