## Supplementary material for "BAllC and BAllCools: Efficient Formatting and Operating for Single-Cell DNA Methylation Data": tabS1.ballc-meta-format.pdf

| Index | Field name | Example | Note |
|---|---|---|---|
| 1 | ref | chr12 | |
| 2 | pos | 18283342 | 1-based |
| 3 | strandness | + | either + or - |
| 4 | c_context | CGT | 3 bases starting with “C”; can use bases defined by IUPAC (eg. H=A,T,orC, N=A,T,C, or G, etc) |
