## Supplementary figures and images for "BAllC and BAllCools: Efficient Formatting and Operating for Single-Cell DNA Methylation Data"

### figS1.merging-time.pdf

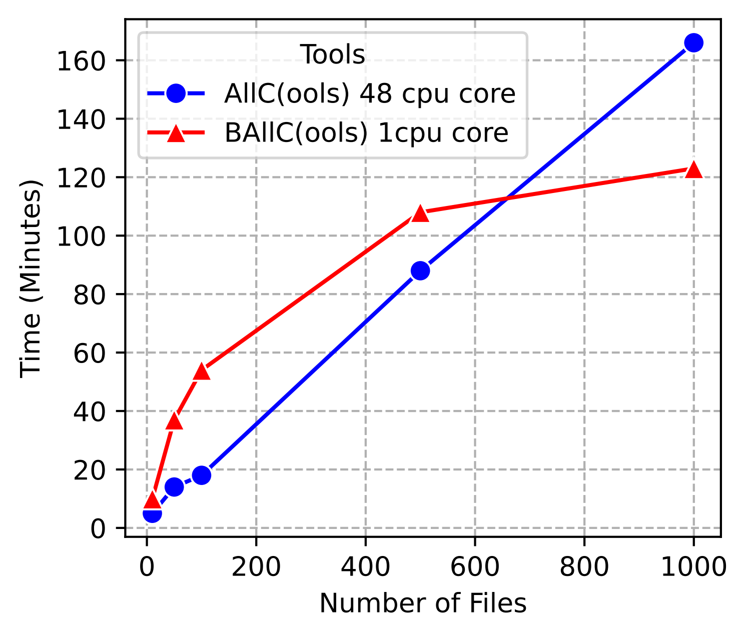
